# Cancer resistance to therapy by tissue-level homeostatic feedback

**DOI:** 10.64898/2026.03.25.714177

**Authors:** Jonathan Somer, Ravid Straussman, Uri Alon, Shie Mannor

## Abstract

Cancer displays remarkable robustness, exemplified by its ability to develop resistance to virtually every therapy. Resistance has traditionally been explained by clonal selection of pre-existing mutations, but there is now abundant evidence for resistance by non-genetic pathways including signals from normal stromal and immune cells. It is largely unclear why normal cells help cancer cells overcome treatment. We propose that physiological circuits responsible for tissue homeostasis can explain why cells cooperate to produce pathological resistance to therapy. To show this, we construct mathematical models of physiological dynamics. We then simulate cancer treatments within the context of a functioning tissue. We find that classic examples of resistance to therapy can be explained by homeostatic feedback regulation - including BRAF inhibitors in melanoma and anti-angiogenic therapy. The *homeostatic theory of resistance* (HTOR) reframes resistance as a byproduct of tissue robustness, rather than solely tumor-specific adaptation. Finally, we analyze two large-scale single-cell RNAseq databases of normal and cancer samples: the Tabula Sapiens^1^ and the Curated Cancer Atlas^2^. We show that in multiple cancers (breast, colon, kidney, liver, lung, ovary, prostate, and skin), malignant cells preserve their tissue-specific homeostatic cell-signaling. We thus expect the robust feedback loops from healthy tissues to play a role in cancer.

## Introduction

Homeostasis is the primary framework for explaining robustness of biological systems. At the systemic level, organisms maintain body temperature and blood pressure, regulate blood levels of CO_2_, glucose and calcium^3^. At the tissue level, organs and bones maintain a stable mass and many tissues recover from injury^4^. Cells regulate their membrane potential and concentrations of numerous metabolites^5^.

This robustness is universally achieved through feedback regulation, similar to engineered control systems^6–9^ (Fig 1a, left). For example, high blood glucose increases pancreas insulin secretion, which activates processes that lower glucose levels back to normal^3^ (Fig 1a, right).

**Figure 1:**
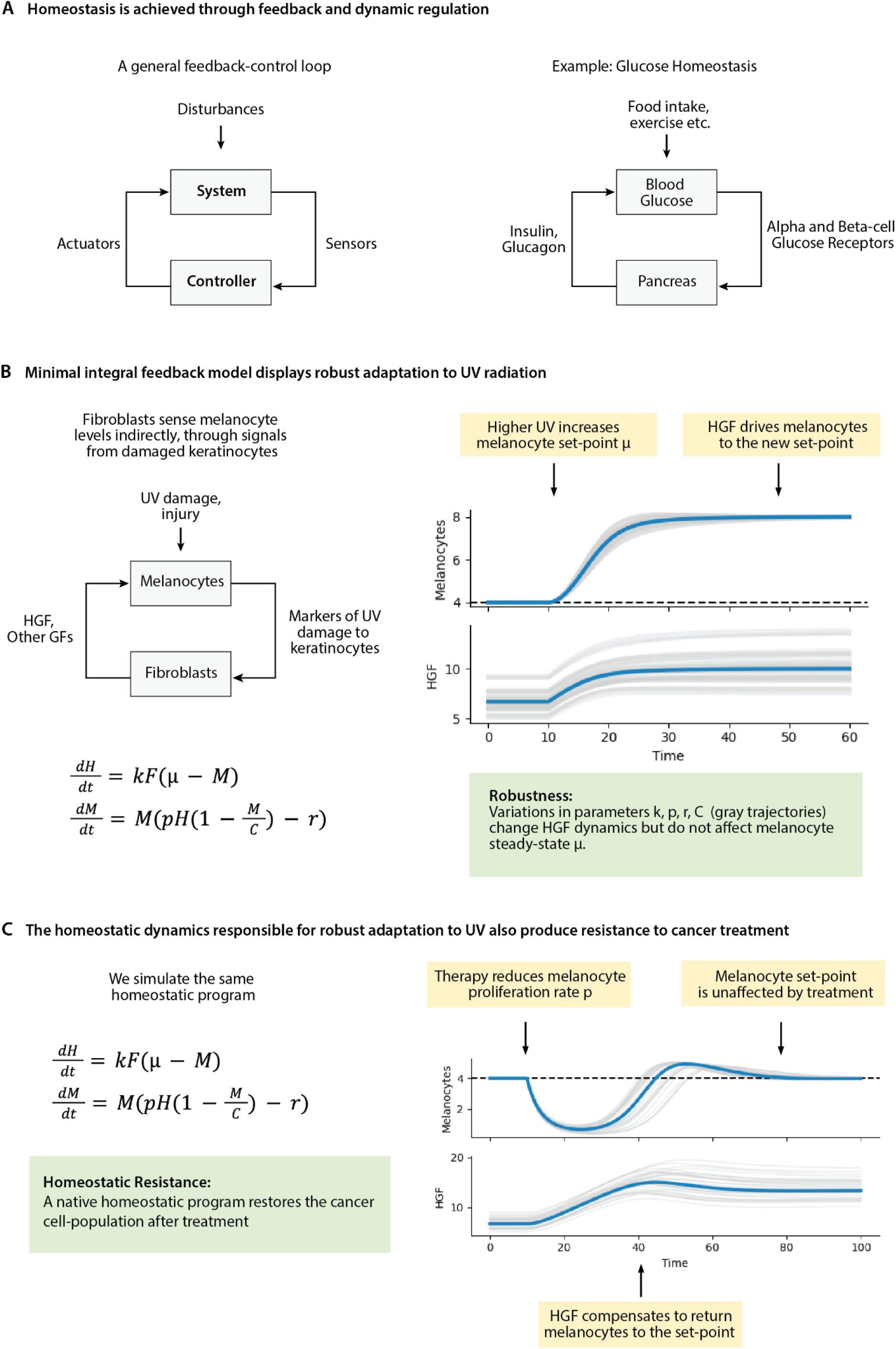
Feedback-control underlies tissue homeostasis and offers a non-mutational mechanism for cancer resistance to treatment. A) Homeostasis is achieved through feedback and dynamic regulation. A controller senses the system’s state and regulates it using actuators (left). Glucose homeostasis can be viewed as a feedback-control loop where the pancreas is the controller (right). B) A minimal integral-feedback model displays robust protection from UV. In this model fibroblasts are viewed as a controller of melanocytes. C) The same integral-feedback model produces resistance to therapy. Therapy is modeled as a 10x reduction in melanocyte proliferation rate.

Cancer’s ability to recover from treatment and acquire resistance is also a form of robustness ^10–12^. Traditionally resistance was explained by clonal selection of resistant cells^13^. In human subjects, the strongest evidence for mutation-driven resistance comes from mutant-targeted treatments, such as BRAF^V600E^ inhibitors in melanoma^14^. Melanoma patients carrying the mutation initially respond to treatment, but disease typically progresses within 5-8 months^14^. Mutations associated with resistance are found in over 50% of patients, but a considerable fraction of resistance cases remain unexplained^14^. An even larger fraction remains unexplained in treatments that do not target specific mutations, such as chemotherapy or immunotherapy^15–18^.

Unexplained cases of resistance may be driven by mutations that we haven’t found, but there is accumulating evidence of non-genetic drivers of resistance^15,16,19,20^. In addition, non-genetic mechanisms may produce conditions that enable mutations to induce resistance.

Important non-genetic drivers of resistance include stromal and immune cells within the tumor-microenvironment (TME) ^19–24^. These normal cells often respond to therapy in ways that help the cancer cell population. For example, in melanoma patients treated with BRAF inhibitors, fibroblasts upregulate their secretion of HGF, a growth factor that makes melanoma cells resistant to BRAF inhibitors^20^.

Why does a normal fibroblast release HGF following a drug that specifically targets mutant-BRAF within melanocytes? More broadly, do tissues have some innate organization that enables cancers to hijack normal cells to support their own survival in the face of treatment?

We propose that native homeostatic feedback loops can explain why cancer cells and stromal cells cooperate to overcome treatment. We propose a theory and mathematical framework for explaining the role of normal cells in cancer resistance to therapy. This theory also sheds light on why specific mutations tend to appear in particular tissues^25^ and why some changes in local or systemic physiology support the tumor.

Our theory is grounded in the mathematical definition of *adaptation*^7,26–28^. A dynamical system is well-adapted if a large change in input produces just a small change in output. In the examples above, the natural outputs are the targets of homeostasis (e.g blood glucose or body temperature). The input could be any external disturbance, such as a meal or a change in ambient temperature.

Importantly, the same system may be well-adapted with respect to multiple input-output pairs. We propose that tissues are expected to be well-adapted to cancer treatments as a side-effect of their ability to robustly perform their function.

To show this we construct mathematical models of tissue physiology based on clinical and experimental data. We then simulate perturbations that mimic cancer treatments within the native physiological context of a tissue. These simulations expose innate compensatory mechanisms capable of rescuing the tumor from treatment.

We model BRAF inhibitors in melanoma, and link resistance to the homeostatic role of melanocytes - protection of skin from UV. We also model anti-angiogenesis therapy, and link resistance to native homeostasis of tissue-oxygenation. In both cases the same dynamics responsible for normal tissue function also produce resistance to therapy - without mutations or selection.

Finally, we analyze two large-scale single-cell RNAseq databases and show that across multiple common malignancies, cancer cells retain the characteristic cell-cell signaling pathways from their healthy cells-of-origin.

### A minimal model of tissue homeostasis displays resistance to therapy

To illustrate how a homeostatic program can produce resistance to cancer therapy we develop a minimal mathematical model of skin UV-damage homeostasis – the tanning response.

The primary skin cell is the keratinocyte. Keratinocytes are protected from UV by cells known as melanocytes^29–31^. Melanocytes produce the protective pigment melanin, and deliver it to keratinocytes. Melanocytes are also the cell of origin in melanoma, the most aggressive skin cancer^32^. When keratinocytes experience UV damage they signal that more melanin and melanocytes are needed ^33–35^. Fibroblasts respond to these signals by secreting growth factors such as HGF, which increase melanocyte proliferation^30,36,37^.

In our minimal model fibroblasts secrete HGF as a response to deviation of melanocyte density from a set-point μ, representing an optimal density of melanocytes under some fixed level of UV. We assume melanocytes have a constant removal rate and a maximal carrying capacity (Fig 1b, left).

In this feedback loop fibroblasts are a controller of melanocytes (Fig 1b, left). If a change in UV intensity shifts the melanocyte set-point μ or if melanocyte levels drop due to injury, fibroblasts adapt their HGF production and drive melanocytes back to the set point (Fig 1b, right).

In this model, fibroblasts regulate melanocytes through integral feedback. This form of control is *robust* because melanocyte steady-state levels don’t depend on any model parameters except the set-point μ^7,27,38–40^. Thus, at steady-state we are guaranteed to have sufficient UV protection (Fig 1b, right).

To see how native homeostatic dynamics can produce resistance, we simulate a cancer therapy that reduces melanocyte proliferation by 10x. We keep all other model parameters fixed (Fig 1c, left). The perturbation mimics a continuously administered drug. Following the perturbation, melanocyte levels transiently drop. HGF then rises to compensate and melanocytes return to pre-treatment levels.

In all respects, this appears as an acquired resistance to therapy – we continuously administer the drug and it eventually stops working. Yet no mutations or selection were present in the process. We also explain the rise in fibroblast HGF following a treatment that specifically targeted melanocytes.

### The UV damage homeostasis circuit explains melanoma resistance to BRAF inhibitors

The minimal model illustrated the core argument with minimal assumptions. We now present a more detailed model grounded in clinical and experimental studies of UV homeostasis. We show that resistance through homeostatic feedback occurs when we account for the signaling pathways, timescales, and regulatory architecture of the skin (Figure 2A).

**Figure 2:**
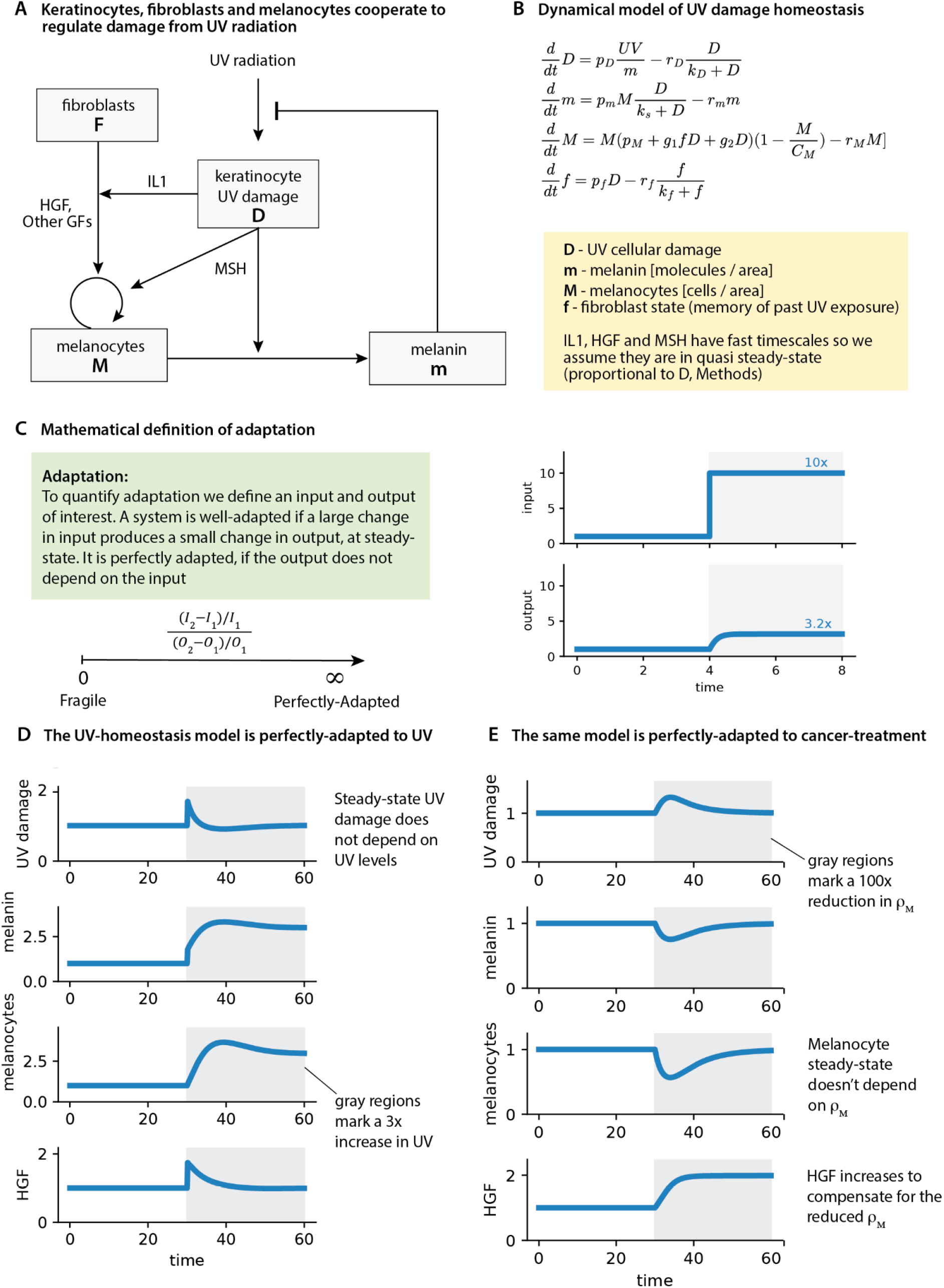
UV damage homeostasis explains melanoma resistance to BRAF inhibitors. A) Keratinocytes, fibroblasts and melanocytes cooperate to regulate damage from UV radiation. B) Dynamical model of UV damage homeostasis. C) Mathematical definition of adaptation. D) The UV-homeostasis model is perfectly-adapted to UV. E) The same model is also perfectly-adapted to cancer-treatment (reduction in *p*_*M*_).

We begin by modeling UV damage (*D*) in keratinocytes. Many forms of DNA damage are produced proportionally to *UV* intensity^41^ and accumulate inversely with melanin content^42^. Following UV exposure, low levels of DNA damage are typically repaired within 24 hours^43^.

When keratinocytes are damaged, they produce signals such as MSH which drive melanocytes to proliferate^44^ and produce more melanin^45^. They also influence melanocytes indirectly, by signaling to fibroblasts via factors such as IL-1^36^. IL-1 increases fibroblast production of growth factors such as HGF^36,37,46^. Timescales of MSH, IL-1, and HGF are fast relative to changes in melanin and melanocytes (Methods, S4). We therefore assume they are in quasi steady-state, which we show is proportional to UV damage *D* (Methods).

In addition to the immediate production of growth factors, fibroblasts also undergo slower, lasting changes in response to UV^47–51^. A single UV exposure causes fibroblasts to gradually upregulate HGF over several weeks^51^. With cumulative UV exposure, fibroblasts progressively acquire a melanogenic phenotype. This process is known as photo-aging, and it underlies various age-related pigmentation disorders^47–51^.

Thus, UV induces both immediate and long-lasting changes in the tissue. The memory of previous UV exposure is essential for keeping the system calibrated to its environment — for instance, by expanding the melanocyte population to maintain a margin for future exposures to UV.

We checked that the model (Figure 2B) performs the primary function of this homeostatic circuit - adaptation to UV. A system is adapted if a large change in input produces just a small change in output, at steady state (Figure 2C, Methods). A system is *perfectly adapted*, if the output does not depend on the input at all.

The UV homeostasis model exhibits perfect adaptation. We simulate a 3x increase in UV intensity, and after a transient period of adjustment damage (*D*) returns to its original level (Figure 2D). Perfect adaptation holds in general — the analytically derived steady-state expression for damage does not include a term for UV (Methods, Table S3).

We now change our perspective and consider the melanocytes as the output and cancer-treatment as the input. Melanoma driven by the BRAF^V600E^ mutation is often treated with BRAF inhibitors. These targeted drugs block the constitutively active pathway caused by the mutation. We thus model treatment as a reduction in the parameter *p*_*M*_ (melanocyte constitutive proliferation rate).

We find that the same model is perfectly-adapted to cancer treatment (Figure 2E). As before, the steady-state formula for Melanocytes (*M*) does not include a term for *p*_*M*_. In fact, melanocyte steady-state does not depend on any of the melanocyte model parameters (*p*_*M*_, *r*_*M*_, *g*_2_, *g*_1_ ). From an evolutionary perspective, this is required. Melanocytes are long-lived cells that are constantly exposed to UV. As such, they may acquire alterations that modify their turnover rate or response to external factors. The tissue must maintain its function despite these alterations.

In summary, this circuit evolved to robustly regulate skin UV damage. As a result it is robust to variations in melanocyte parameters. Treatments that modify melanocyte parameters are thus counteracted by native compensatory feedback loops.

### Tissue oxygen homeostasis explains resistance to anti-angiogenic therapy

The previous example focused on a treatment that targets mutant BRAF within cancer cells. We now turn to anti-angiogenesis therapy, which targets a process external to the cancer cells.

Angiogenesis, the growth of blood vessels, is a hallmark of cancer^52^ and has been considered a promising therapeutic target^53^. The pioneering anti-angiogenic drugs target the growth factor VEGF or its receptor. Anti-angiogenic drugs were approved for a number of cancers, yet in most cancers improvements in progression-free survival remain on the order of a few months^54^. Resistance remains a key challenge.

A common mode of resistance to anti-angiogenic therapy is upregulation of compensatory pro-angiogenic factors by tumor and stromal cells^55–58^. Following VEGF-blockade, angiogenic factors that played a minor role before treatment suddenly rise and become drivers of angiogenesis. Compensatory factors include FGF, angiopoietin and ephrin^55^. To explain the rise in compensatory factors in the tumor we again turn to the homeostatic dynamics of the tissue.

The primary homeostatic role of angiogenesis is regulating tissue oxygen levels. Nearly every cell in the body can sense and respond to hypoxia^59^. Cells constantly produce the transcription factor HIF1a, and in normal oxygen levels it is rapidly degraded. When oxygen levels drop, HIF1a accumulates and enters the nucleus where it activates hypoxia-related genes and expression of angiogenic factors such as VEGF. Thus, various cell types independently act to correct oxygen levels (Fig 3a).

**Figure 3:**
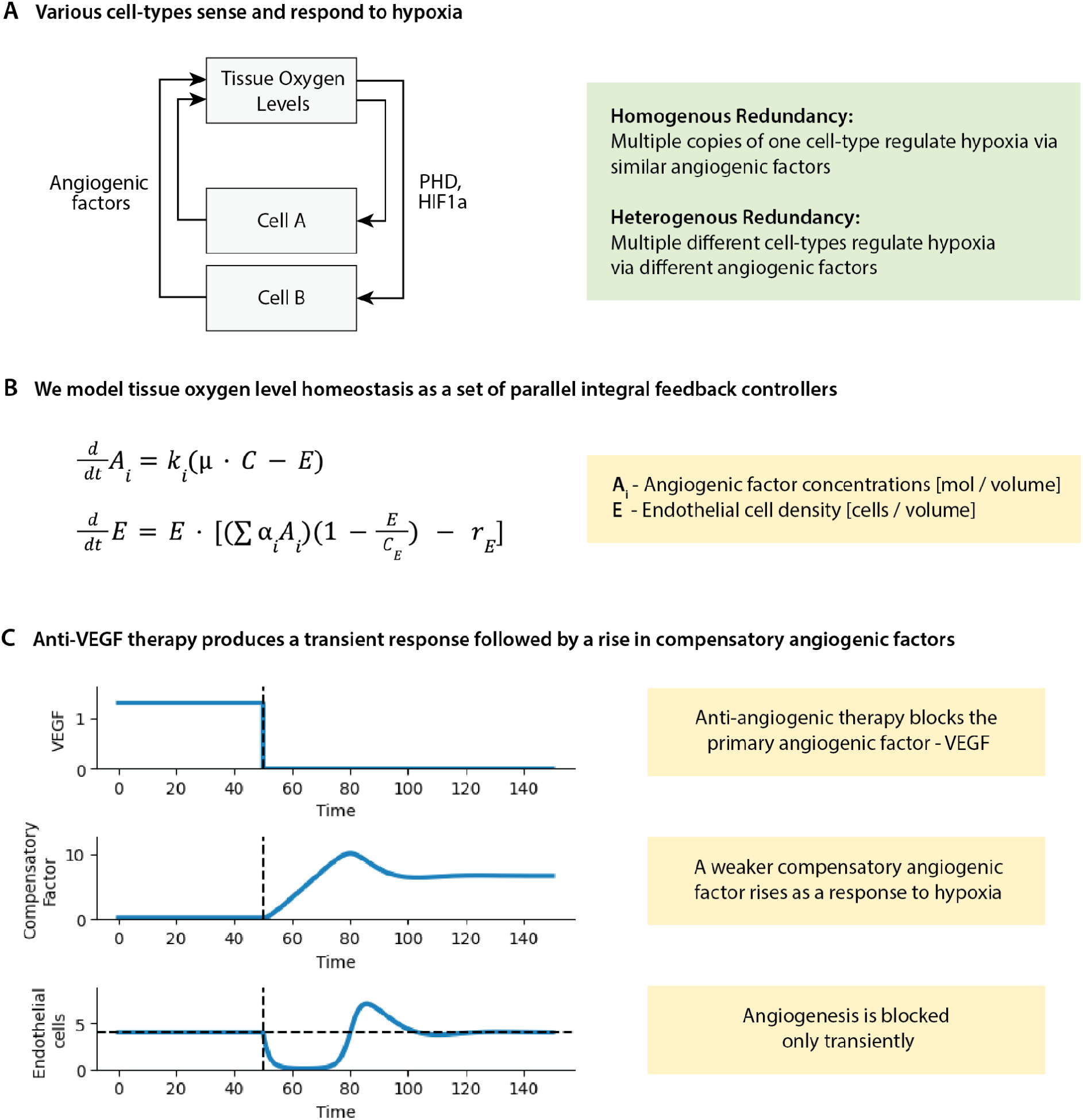
Tissue hypoxia homeostasis explains resistance to anti-angiogenic therapy. A) Various cell-types sense and respond to hypoxia. This feedback-control pattern displays both homogenous and heterogeneous redundancy: multiple copies of each cell-type and various different cell-types. B) We model tissue oxygen homeostasis as a set of parallel integral feedback controllers, each corresponding to a different angiogenic factor. C) Anti-VEGF therapy produces a transient response followed by a rise in compensatory angiogenic factors.

We model this feedback structure as a set of parallel integral feedback controllers, each corresponding with a different angiogenic factor (Fig 3b). We assume normoxia is reached when the ratio of endothelial cells to consumer cells is μ^60^. Cells respond to deviations of endothelial cells from this optimal ratio. For clarity, we simulate two cell types, each secreting a different angiogenic factor, but our results are easily extended to multiple cell types and overlapping secretion of angiogenic factors.

We assume the first cell type responds strongly to hypoxia (large *k*_1_ ) and produces VEGF, which has a potent effect on endothelial cell proliferation (large *a*_1_ ). The second cell-type responds more weakly to hypoxia (small *k*_2_), and produces an angiogenic factor with a smaller effect on endothelial cells (small *a*_2_).

We simulate anti-VEGF therapy by setting VEGF levels to 0. We find that following VEGF blockade, endothelial cells transiently drop, followed by a rise in the weaker growth factor (Fig 3c). The weaker growth factor had a negligible effect on endothelial cell proliferation before VEGF blockade, but now it is the primary driver of angiogenesis.

Once more, this appears as an acquired resistance to therapy - we continuously administer the drug and it eventually stops working.

### Mutations and anomalous physiological contexts can modify native homeostatic feedback loops

The *homeostatic theory of resistance* (HTOR) is compatible with the clonal theory of resistance. In fact, homeostatic feedback loops may be necessary for mutations to produce resistance. For example, following treatment, compensatory feedback loops may be necessary for cancer cells to survive long enough to acquire resistance mutations.

In addition, homeostatic feedback loops may create points of leverage that can be exploited by mutations. For example, based on the UV homeostasis model (Fig 2a) we expect melanocytes to benefit from reduced melanin production, because keratinocytes signal that more melanocytes are needed when they aren’t sufficiently protected from UV (Fig 4a).

**Figure 4:**
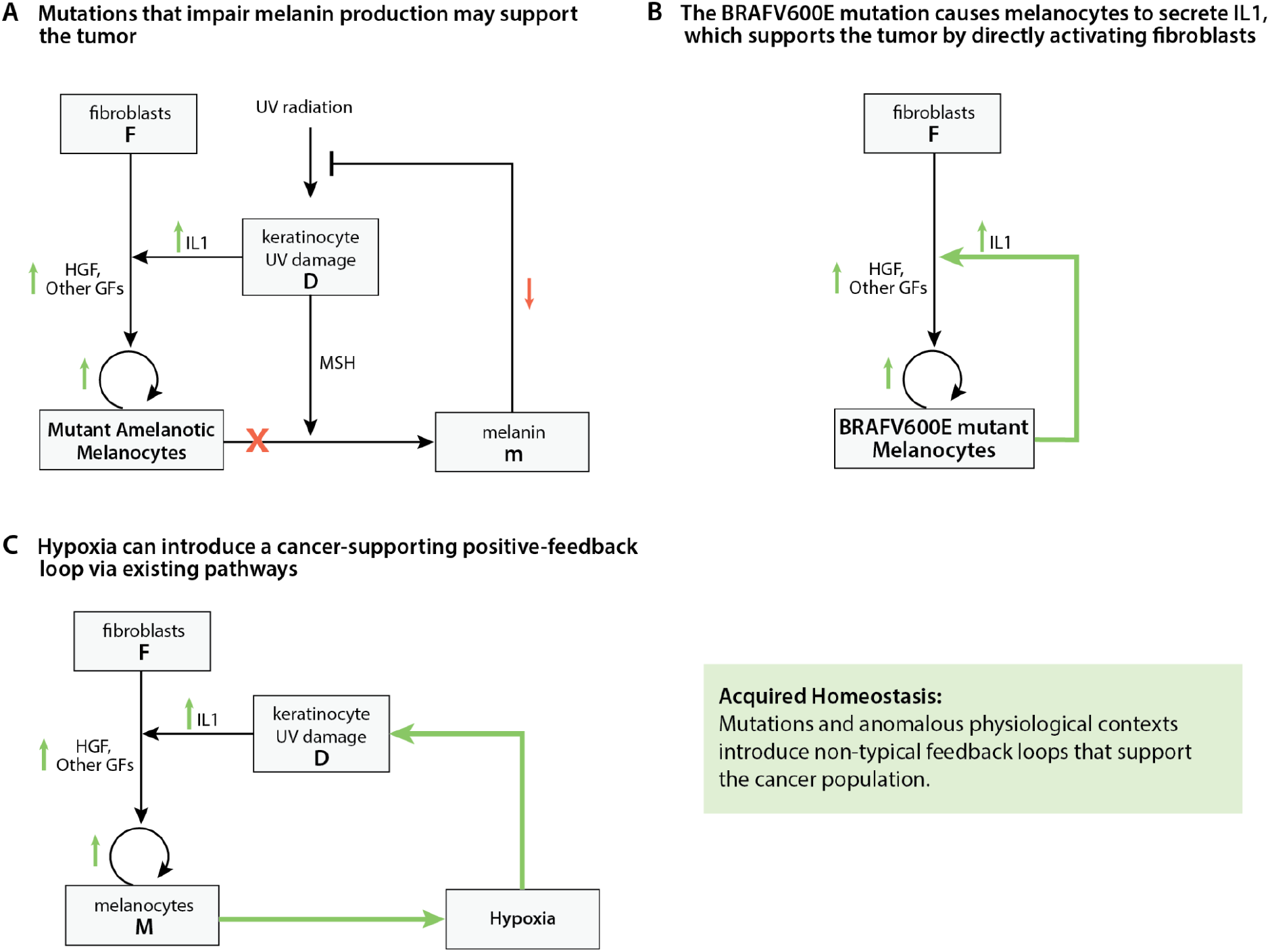
Mutations and anomalous physiological contexts can modify native homeostatic feedback loops. A) Mutations that impair melanin production may support the tumor. B) The BRAFV600E mutation causes melanocytes to secrete IL1, which supports the tumor by directly activating fibroblasts. C) Hypoxia can introduce a cancer-supporting positive-feedback loop via existing pathways.

Indeed, 8% of melanomas do not produce melanin and are known as *Amelanotic Malignant Melanoma* (AMM)^61–63^. AMM is harder to diagnose and has worse prognosis. Notably, AMM also grows faster, as expressed by a higher mitotic index^61^.

It is challenging to explain the rapid proliferation rate in AMM without the physiological role of melanin. The most common mutations responsible for loss of melanin production in AMM are albinism genes TYR and OCA2^64^. These genes have specific roles in melanin secretion and to the best of our knowledge have no established intracellular role in reducing melanocyte proliferation^62^.

Based on our model, IL1 overexpression should also support melanocytes (Fig 4b). Remarkably, the BRAF^V600E^ mutation, which drives 50% of melanoma cases, also induces IL1 expression by melanocytes and activates adjacent fibroblasts as predicted by the homeostatic model^65^. This is a strong example of how the homeostatic and genetic perspectives complement each other.

Besides mutations, anomalous physiological contexts may also perturb native homeostatic feedback loops. For example, melanomas are typically hypoxic^66^, as are the cores of most solid tumors. Hypoxia produces reactive oxygen species and triggers IL1 production by keratinocytes^67^. This introduces a positive feedback loop that supports the cancer - proliferating melanoma cells induce hypoxia, this increases IL1-HGF signaling, which further increases melanoma cell proliferation (Fig 4c). Similarly, systemic physiology may also modify tissue-level homeostatic circuits to the tumor’s advantage^22^.

### Pan-cancer single-cell RNAseq data shows homeostatic cell-signaling is preserved in cancer

The homeostatic theory of resistance (HTOR) is rooted in the robustness of homeostatic tissue physiology. A key assumption is that cancer cells retain at least some of the homeostatic circuitry of their healthy predecessors. This preservation is in contrast to the well-known changes in cancer metabolism and loss of effector functions (e.g. thyroid hormone production in thyroid cancer and enzyme production in liver cancer).

If indeed cancer cells sense and respond to similar homeostatic signals as the normal cells from which they originated, we can expect the robust feedback loops from the healthy tissue to play a role in cancer.

The literature on cancer often emphasizes aspects that are unique to the malignant tissue. Cancer cells are depicted as mutated, transformed or de-differentiated. In the following analysis, we set out to test if cancer cells retain the cell-cell interactions from the healthy tissue. Namely, do the inputs (receptors) and outputs (ligands) of cancer cells resemble those of their corresponding cells-of-origin?

It is important to consider both inputs and outputs, because a cell could express a receptor but respond differently, due to downstream alterations. Preservation of both receptor and ligand profiles suggests that the core signaling logic — from input to output — remains largely intact, even if individual downstream pathways are modified.

We aim to detect similarities in cell-cell interactions between cancers and the normal tissues they developed from. During analysis, such similarities could potentially arise from experimental batch effects or artifacts in data preprocessing. That is, if normal and tumor samples are taken from the same studies, then similarities in gene expression may be attributable to the technical factors above. To overcome this potential pitfall, we took normal and cancer samples from entirely different studies. Thus, all inferred similarities were robust to differences across both patients and experimental batches.

We analyzed two large-scale single-cell RNA-seq databases. We took cancer samples from the curated cancer atlas^2^, which includes a total of 2836 samples from 124 studies. We used normal samples from the Tabula Sapiens database which includes normal tissue samples of various organs from 15 donors^1^.

We selected 8 organs from the Tabula Sapiens database - breast, colon, kidney, liver, lung, ovary, prostate and skin. Multiple types and subtypes of malignancies can develop from each organ. To simplify the analysis, we chose the malignancy with most samples in the curated cancer atlas (Table S7). We did not distinguish between cancer subtypes such as (HER-2 positive or triple-negative breast cancer).

We focus on malignant cells from tumor samples and epithelial cells from normal samples. In normal samples, we included only epithelial cells that are considered as the cell-of-origin for the corresponding malignancy (Table S7). To avoid issues related to varying statistical power when comparing samples with different numbers of cells, we worked with pseudobulk samples (Methods).

We computed spearman rank correlations of ligand and receptor gene expression levels, between each normal and each cancer sample, across all organs. We find that cancer cells are most similar to epithelial cells from the same tissue (red diagonal blocks in Figure 5A).

**Figure 5.**
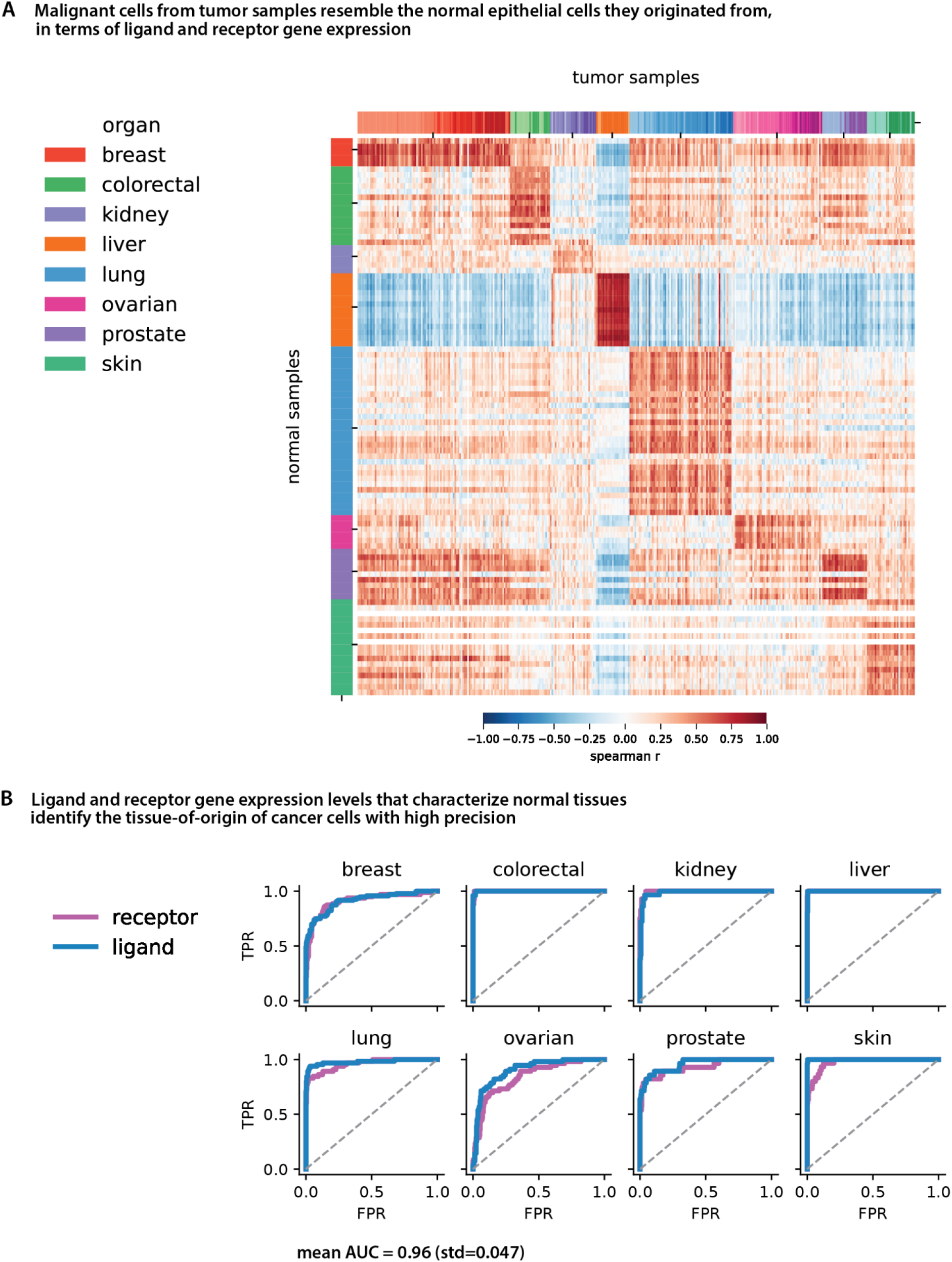
Pan-cancer single-cell RNAseq data shows cancer cells preserve their tissue-specific homeostatic cell-signaling. A) Spearman rank-correlation between pseudobulk ligand and receptor gene expression in epithelial cells from normal and cancer samples. Data includes 99 normal samples and 350 tumor samples. Red blocks on the diagonal imply malignant cells from tumor samples resemble the normal epithelial cells they originated from, in terms of ligand and receptor gene expression. B) Ligand and receptor gene expression levels that characterize normal tissues identify the tissue-of-origin of cancer cells with high precision. We fit logistic regression models to ligand or receptor gene expression data (quantiles), with the aim of detecting the organ an epithelial cell is from (Methods). The model was fit only to data from healthy tissue samples. Each panel depicts receiver operating characteristic (ROC) curves for classification of one tissue type, based on ligand or receptor gene expression in malignant cells. The model detects the tissue type with high precision (AUC mean: 0.96, std: 0.047, Table S3).

To further test the extent of cell-signaling preservation, we fit logistic regression models to ligand or receptor gene expression data (quantiles), with the aim of detecting the organ of origin of an epithelial cell (Methods). The model was fit only to data from healthy tissue samples. In order to distinguish between an epithelial cell from breast vs colon, the model must pick up on receptors (inputs) or ligands (outputs) unique to each tissue. We inspected the 10 largest coefficients for each tissue (Additional material, Tables S9,10). As expected, the model picks up on genes with canonical homeostatic roles. Examples include the prolactin receptor in breast tissue (lactation), the angiotensin-2 receptor in lung tissue (blood pressure and fluid homeostasis) and parathyroid hormone receptor in kidney tissue (calcium homeostasis).

As predicted by HTOR, the models generalize to cancer samples without any modification (Figure 5B). They detect the tissue type of malignant cells with high precision, based on similarity of their ligands or receptors to those of healthy epithelial cells from each tissue (mean AUC: 0.96, std: 0.047, S8). We conclude that cell-cell interactions involved in homeostatic tissue function are preserved in malignant cells.

## Discussion

We propose that tissue-level homeostatic feedback may be sufficient to generate resistance to therapy—without genetic mutations or clonal selection. By modeling the physiological dynamics of tissues and introducing perturbations analogous to cancer therapies, we show that resistance can emerge naturally from the same regulatory circuits that evolved to maintain healthy tissue function. This framework, which we term the *homeostatic theory of resistance* (HTOR), offers a unified explanation for diverse resistance phenomena and provides a quantitative method to predict which compensatory mechanisms a given treatment is likely to trigger.

Our results support the view that resistance is not solely a property of cancer cells, but can be an emergent property of the tissue control system. The approach also offers mechanistic context for why certain mutations—those that disrupt effector functions or hijack existing feedback loops—are particularly advantageous for tumor progression and therapy evasion.

We examined two broad classes of treatments - treatments that target cancer cells and treatments that target processes external to cancer cells. From the first class, we modeled resistance to BRAF inhibitors in melanoma within the context of skin UV-damage homeostasis. This model explains the otherwise puzzling induction of fibroblast-derived HGF following melanocyte-specific inhibition. It also clarifies why mutations that impair melanin production or induce IL1 secretion can accelerate disease progression.

Because many cancers originate from cells with distinct physiological roles, we hypothesize that the resistance pathways in these cancers should follow similar patterns to those we found in melanoma. Namely, loss of specific effector functions (such as melanin) coupled with stromal upregulation of trophic factors (such as fibroblast-derived HGF). In some cases, tumor cells may directly activate stromal cells via canonical physiological pathways (such as melanocyte derived IL1, which mimics keratinocyte UV-damage signals).

We also considered therapies that target processes external to cancer cells. We modeled resistance to anti-angiogenic therapy by simulating VEGF blockade in the context of tissue oxygen homeostasis. We identified a robust parallel-control structure in which multiple cell types independently sense hypoxia and secrete angiogenic factors. In this architecture, inhibiting one factor inevitably leads to compensatory upregulation of others.

We expect this control motif to underlie other cases of unexpected compensatory signaling, especially in fundamental homeostatic processes such as inflammation or immune regulation. For example, various cell types can sense inflammation and produce immunosuppressive signals. Thus, we predict that immune checkpoint blockade which typically inhibits one immunosuppressive pathway, should inadvertently increase activity in other pathways. A similar result was shown by Koyama et. al, where T-cells upregulate alternative immune checkpoints after PD-1 blockade^68^.

The HTOR offers an explanation for transient resistance^18,69,70^ - breaks from treatment provide opportunity for tumor-supporting feedback to decay. Timing treatments based on markers of adaptation could improve efficacy over protocols that resume treatment only upon disease progression. We develop such strategies in detail in a forthcoming publication.

Homeostatic feedback loops provide points of leverage where mutations could have a large effect on the system. They may also provide points of leverage for synthetic feedback elements to steer the system towards tumor suppression. Advances in synthetic biology may make such “homeostasis hacking” feasible^27^.

In sum, it is difficult to treat cancer not only because of its genetic heterogeneity, but because it is embedded within a physiological system that evolved to be robust over millions of years. An important implication of the HTOR is that we should generally expect fixed treatments to fail to achieve a lasting response - robust physiological feedback loops will bypass the perturbation and maintain stability.

## Methods

### Modeling UV homeostasis

Keratinocytes, fibroblasts and melanocytes cooperate to protect skin from UV radiation^29–31^. Keratinocytes are the most abundant and most exposed skin cell^71^. Melanocytes protect keratinocytes by producing melanin.

We start by modeling dynamics of UV damage in keratinocytes. The change in damage is given by the rate of damage production minus rate of damage repair. UV produces damage in a dose-dependent manner (both duration and intensity). For example, many types of DNA damage accumulate linearly with UV dose in isolated DNA samples^41^. DNA damage accumulates inversely with melanin content^42^. Following a single exposure, low levels of DNA damage are repaired within 24 hours43. Some high exposures take much longer to repair, consistent with saturation of the repair machinery^43^. In this model we do not distinguish between different forms of DNA damage, or damage to other cell components. We thus model UV damage dynamics as:

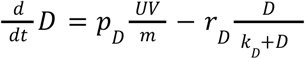

To further simplify analysis, we assume that UV intensities are within the repair capabilities of single cells (*D* << *k*_*D*_ ). Thus, we do not model strategies for handling irreparable damage levels, such as cell-cycle arrest and regulation of keratinocyte turnover, which also play an important role in skin homeostasis.

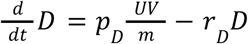

Keratinocytes (*K*) secrete multiple factors in response to UV damage. An inflammasome mediated pathway senses DAMPs (damage associated molecular patterns) and releases IL1 family proteins ^72^. A p53 mediated pathway senses DNA damage and produces MSH^73^. Both pathways are part of the early UV response and have fast kinetics on the order of hours ^34,35^. We assume production rates are hill-function responses to damage and that typical removal rates are on the order of hours. The IL1 model:

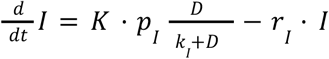

Melanin production and melanocyte proliferation are on the order of weeks and months^31^ so we assume IL1 is in quasi-steady state. As before we assume responses aren’t saturated ( *D* << *k*_*I*_ ).

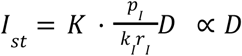

MSH has similar dynamics. Thus, both IL1 and MSH are proportional to damage:

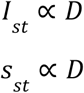

The protective pigment melanin is produced by melanocytes. Melanocytes increase melanin production as a response to pigmentation factors such as MSH (S4)^45^. Melanin removal is primarily driven by keratinocyte turnover which lasts 4-5 weeks^31^. We assume each melanocyte has a maximal melanin production rate *p*_*m*_. In reality, melanocytes do not only produce more melanin, but increase production of a variant that is more protective from UV (eumelanin)^74^. Melanocytes also improve the distribution of melanin to nearby keratinocytes. We assume *p*_*m*_ accounts for these components. Melanin dynamics are then given by:

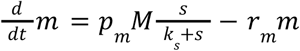

We substitute MSH (*s*) with its steady-state value due to its fast timescale, and assume we are in the unsaturated regime:

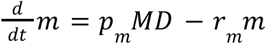

Following a few weeks of UV exposure, melanocyte numbers can increase four-to-sixfold, driven predominantly by cell division^75^. This proliferation is stimulated both directly and indirectly by damaged keratinocytes. Keratinocytes increase melanocyte proliferation directly, via factors such as MSH^44^ (*s*), and indirectly by signaling to fibroblasts through factors such as IL-1^36^. IL-1 increases Fibroblast production of melanogenic growth factors such as HGF (*H*)^36,37,46^. Our model includes terms for both signaling mechanisms, as well as basal melanocyte proliferation and removal rates *p*_*M*_, *r*_*M*_ . We also include a carrying capacity term which limits melanocyte density 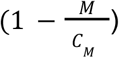.

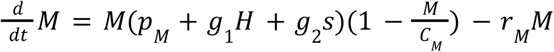

Physiologically, melanocytes are sparsely scattered in the epidermis (a commonly cited number is 1:40 melanocytes to keratinocytes^76^). We therefore assume the density of melanocytes does not directly limit their growth (i.e 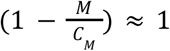).

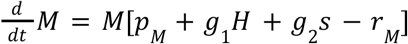

Fibroblasts produce HGF in response to IL-1^36^. HGF approaches steady-state within ∼24 hours (S4)^36^. In addition to this immediate response, fibroblasts also undergo slower and more persistent changes^47–51^. A single UV exposure produces a gradual upregulation of fibroblasts HGF over several weeks^51^. These slow timescale effects occur in parallel to the fast response, and are collectively known as photo-aging. Photo-aged fibroblasts contribute to age-related skin-pigmentation disorders by secreting high levels of melanogenic factors such as HGF^48,50,51^. The process of photo-aging is at least in part driven by accumulation of epigenetic modifications^49,77^. An elegant experiment by Duval et. al^47^ shows that fibroblasts taken from photo-aged skin, can change the pigmentation of reconstructed skin models at the macroscopic level. Thus, fibroblasts harbor memory of previous exposure, independent of current UV levels. We thus model HGF production and removal on a fast timescale, and model the slow timescale changes in fibroblasts as a fibroblast-state variable *f*:

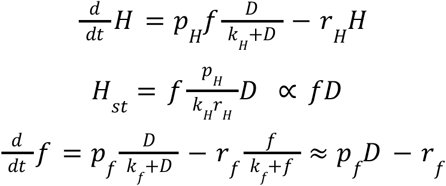

Again, we replace IL1 with its steady-state value *D*. We assume *f* >> *k*_*f*_ so that the state variable integrates previous UV exposure. The final model includes damage (D), melanin (m), melanocytes (M), and fibroblast state (*f*)

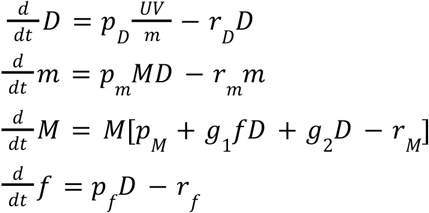

Table S5 summarizes model parameters and units. Table S3 displays analytical formulas of steady-state values, under various alternative modeling assumptions.

### Defining adaptation

Intuitively, a system is well-adapted if a large change in input produces a small change in output, at steady-state. When the output steady-state does not depend on the input at all, we say the system is *perfectly adapted*. “Imperfect” adaptation can be quantified as 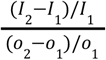. This definition is sensitive to the choice of fold-change in input (*I*_2_, *I*_1_ ), so we propose a definition that is independent of specific values. We define a system as adapted when the output grows sublinearly with the input. When this is the case, 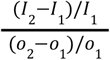 can be made arbitrarily large. For example, let *O*(*I*) = *I*^1−α^, and *I*_2_ = *kI*. Then 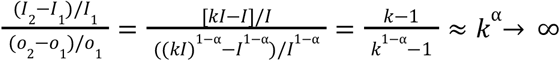 . This definition uniquely determines if an input-output pair is adapted (with the caveat that in practice adaptation may become apparent only for very large inputs).

### UV homeostasis model adaptation

We are interested in two cases of adaptation. The first, is the physiological role of the circuit, namely the change in damage (*D*) as a response to a change in *UV*. The second, is adaptation of the circuit to cancer therapy. In the context of melanoma, parameter *p*_*M*_ (growth-factor-independent proliferation) includes constitutively activated pathways, such as the MAPK/ERK pathway driving BRAF^V600E^ mutant melanoma. We thus model BRAF inhibitors as a reduction in *p*_*M*_ . Therefore, the second type of adaptation we are interested in is the change in melanocyte density (*M*) as a response to change in *p*_*M*_ .

We analytically derive *D*_*st*_ and *M*_*st*_ and consider their dependence on *UV* and *p*_*M*_, respectively. In the case of perfect adaptation *UV, p*_*M*_ do not appear in the formulas at all. In the case of imperfect adaptation, we expect a sublinear term such as 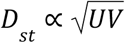.

We derive the steady state formulas under various modeling assumptions (Table S3). We assume melanin production rate is saturated 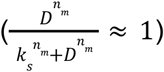 or far from saturation 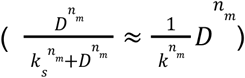 ; we assume melanocyte carrying capacity is a limiting factor (*M* ≈ *C*_*M*_) or not 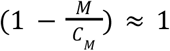; and we assume fibroblasts have long-term adaptation 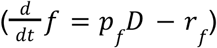 or not. In the latter case, we simplify melanocyte dynamics: 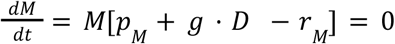.

The complete model displays perfect adaptation in both respects. 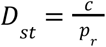 does not depend on *UV*, and 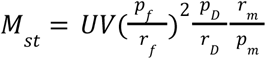 does not depend on *p*_*M*_ . Table S3 shows how the quality of adaptation gradually decreases when different model components are removed or assumed saturated.

### Dimensional analysis of the UV homeostasis model

To identify how model parameters influence the resulting dynamics we employ dimensional analysis. First, we scale *UV, m, M* by reference values 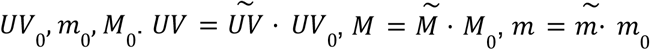. We select a characteristic timescale 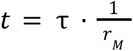. Substituting into the model (removing the ∼ notation for brevity) we get:

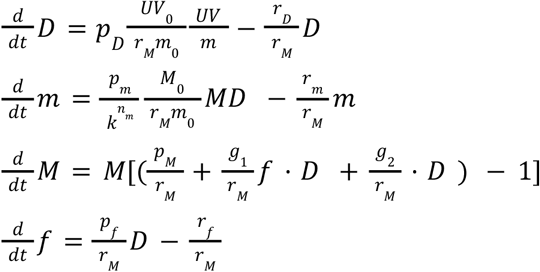

Damage steady-state can be inferred from the equation for 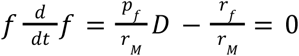, hence: 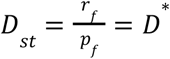. We define 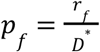 to get 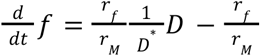. Damage units are arbitrary so we fix the parameter *D*\* = 1. Now we define dimensionless parameters: 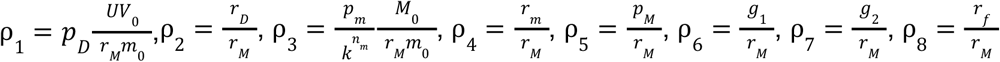.

The dimensionless model is:

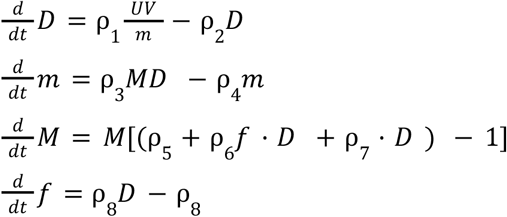

We estimate parameters *r*_*D*_, *r*_*m*_, *r*_*M*_ based on published turnover times (on the order of a day, month, and year, respectively^43,31,78^, Table S1). The turnover time *T* is approximately 5 half lifes, hence the corresponding removal rate is given by 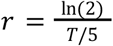. We infer ρ_2_, ρ_4_ using these estimates.

This model displays 3 timescales - fast damage response, intermediate melanin response and slow melanocyte cell population adaptation. We assume fibroblast memory of past exposures has a slow timescale. Specifically, that for every increase in damage on the order of D^*^, a unit of memory is accumulated over the same timescale as melanocyte turnover. 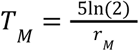 so in the scaled time units, τ_*M*_ = 5 ln(2).

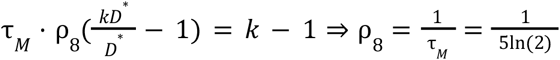

We are left with 5 dimensionless parameters – ρ_1_, ρ_3_, ρ_5_, ρ_6_, ρ_7_. Parameter ρ_1_ defines the rate of damage accumulation for our choice of units *UV*_0_, *m*_0_. Parameter ρ_3_ sets the capacity of melanin production. Parameters ρ_5_, ρ_6_, ρ_7_ tune transient melanocyte dynamics (S1), and do not influence steady-state values:

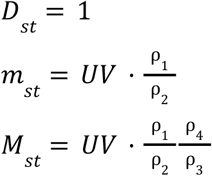

A higher rate of UV production ρ_1_ increases steady-state values of both melanin and melanocytes (S2). Scaling ρ_1_ is equivalent to scaling the units of melanin or UV, so we choose a convenient ρ_1_ = ρ_2_ . Increasing melanin production capacity by melanocytes (ρ_3_) decreases the melanocyte steady-state (S2). We fix ρ_3_ = ρ_4_ such that in our units, *m*_*st*_ = *M*_*st*_ = 1 for *UV* = 1. Figure S2 and table S3 show our conclusions are robust to these choices.

To study the effect of the melanocyte parameters controlling transient dynamics, we first set their values such starting from a steady state where *D*_*st*_ = 1, *f* = 1, each term has equal contribution to melanocyte proliferation rate:

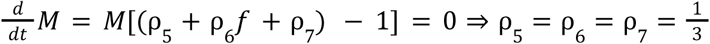. In S1 we study the effect of varying each parameter (S1).

### Single-cell data preparation and exclusions

We performed the following steps:

1. We mapped the granular cell types defined in each study to broad cell types defined in Table S8.
2. We mapped gene names to standard HGNC names using gProfiler.
3. We kept only treatment-naive cancer samples (i.e on-treatment and post-treatment were excluded).
4. We kept only samples collected from the primary tumor site (i.e lymph, blood etc. were excluded)
5. In normal samples we kept only epithelial cells regarded as origin cell types of the corresponding malignancy (Table S7).
6. We performed a quality control step for each study separately:
  a. Remove cells with low complexity (<500 detected genes)
  b. Remove cells with both >10% mitochondrial genes, and <15 housekeeping genes^79^.
7. To generate pseudobulk samples, we summed the raw counts of each cell type.
8. We dropped genes that didn’t have any positive counts across all subjects.

We didn’t normalize or transform the counts data. Instead, we base our analysis on within-sample gene expression ranks or quantiles.

### Computing correlations between ligand and receptor expression in normal vs tumor samples

We constructed a list of all 1655 ligand and receptor genes from the ConnectomeDB 2025 database^80^. To compute correlations we used spearman rank correlations between the pseudobulk counts for each subject. In order to highlight differences in epithelial cells from different tissues, we restricted the analysis to the top 100 most variable genes across all samples. To compute the most variable genes we normalized counts to counts per million and applied the function highly_variable_genes from scanpy (‘seurat_v3’ flavor).

### Detecting the tissue type of cancer cells based on ligand and receptor gene expression

We fit L1-regularized logistic regression models to pseudobulk gene expression from normal samples. We fit one model per organ (e.g. a model that detects breast vs other organs). We fit separate models to ligand and receptor gene expression data. The input to the model was the quantile of each gene within a sample. We tested these models on cancer samples without performing any modifications. We selected the value of the regularization parameter based on a leave-one-out cross validation on normal breast samples, and used the same regularization constant across all models.

## Supplementary Guide

**Figure S1.**
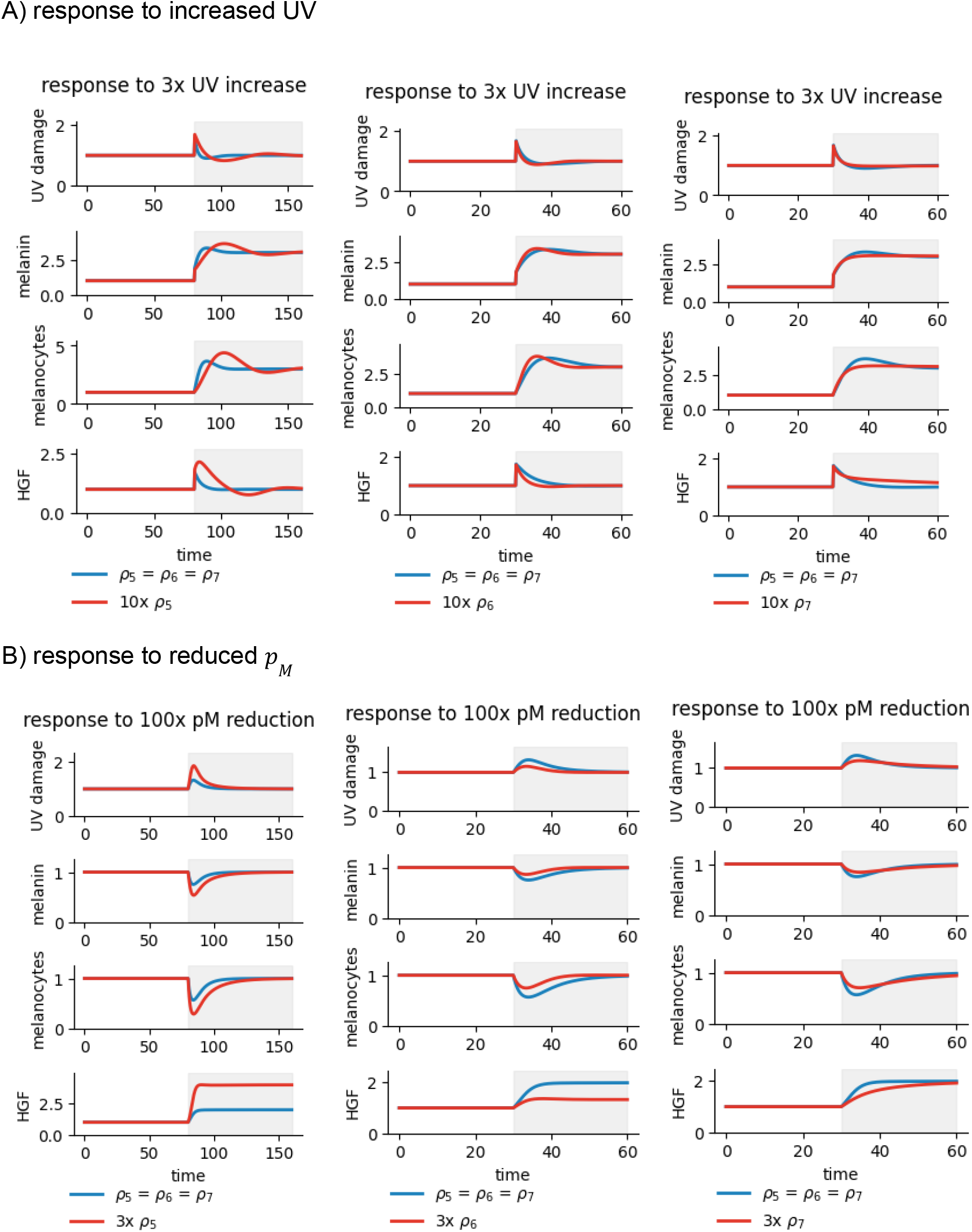
dimensionless parameters ρ_5_, ρ_6_, ρ_7_ tune transient dynamics, but do not influence the steady state. In the main text, we used a model where ρ_5_ = ρ_6_ = ρ_7_ = 1/3 (Methods). Here we plot trajectories where the ratio between parameters is not 1:1:1 but rather 1:1:10 for each parameter at a time. We observe that when ρ_5_ is large relative to ρ_6_, ρ_7_ adaptation takes longer. We also observe that ρ_7_ dampens the response, avoiding over- and under-shoots. A) Response to a 3x increase in UV (gray area). B) Response to a 100x reduction in *p*_*M*_

**Figure S2.**
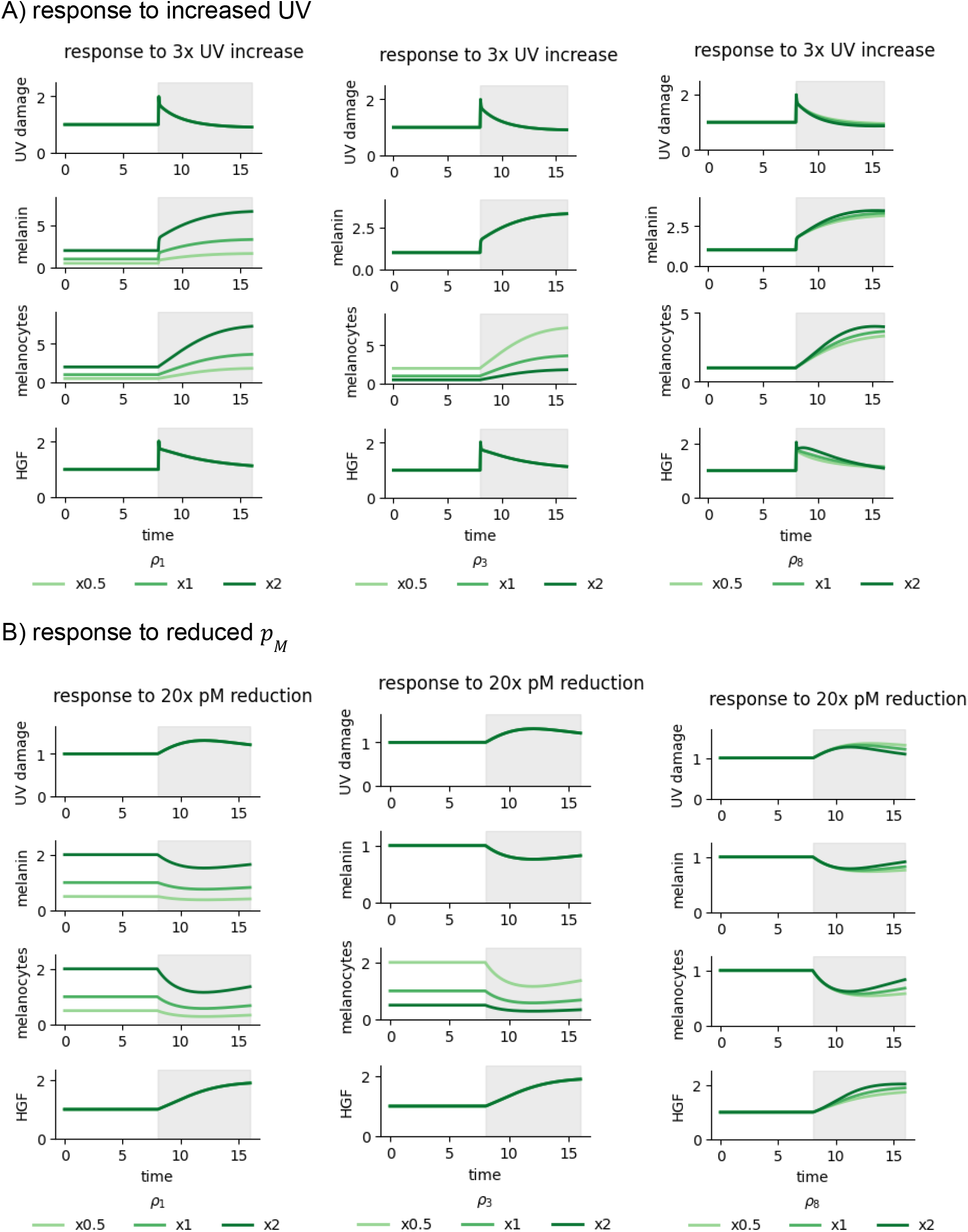
effect of dimensionless parameters ρ_1_, ρ_3_, ρ_8_. Changing parameter ρ_1_ (damage production rate) scales *m, M*. Changing ρ_3_ (melanin production capacity) scales *M* (e.g half the melanin, implies double the melanocytes). ρ_8_ mostly influences long-term adaptation. A) Response to a 3x increase in UV (gray area). B) Response to a 100x reduction in *p*_*M*_.

**Table S3:**
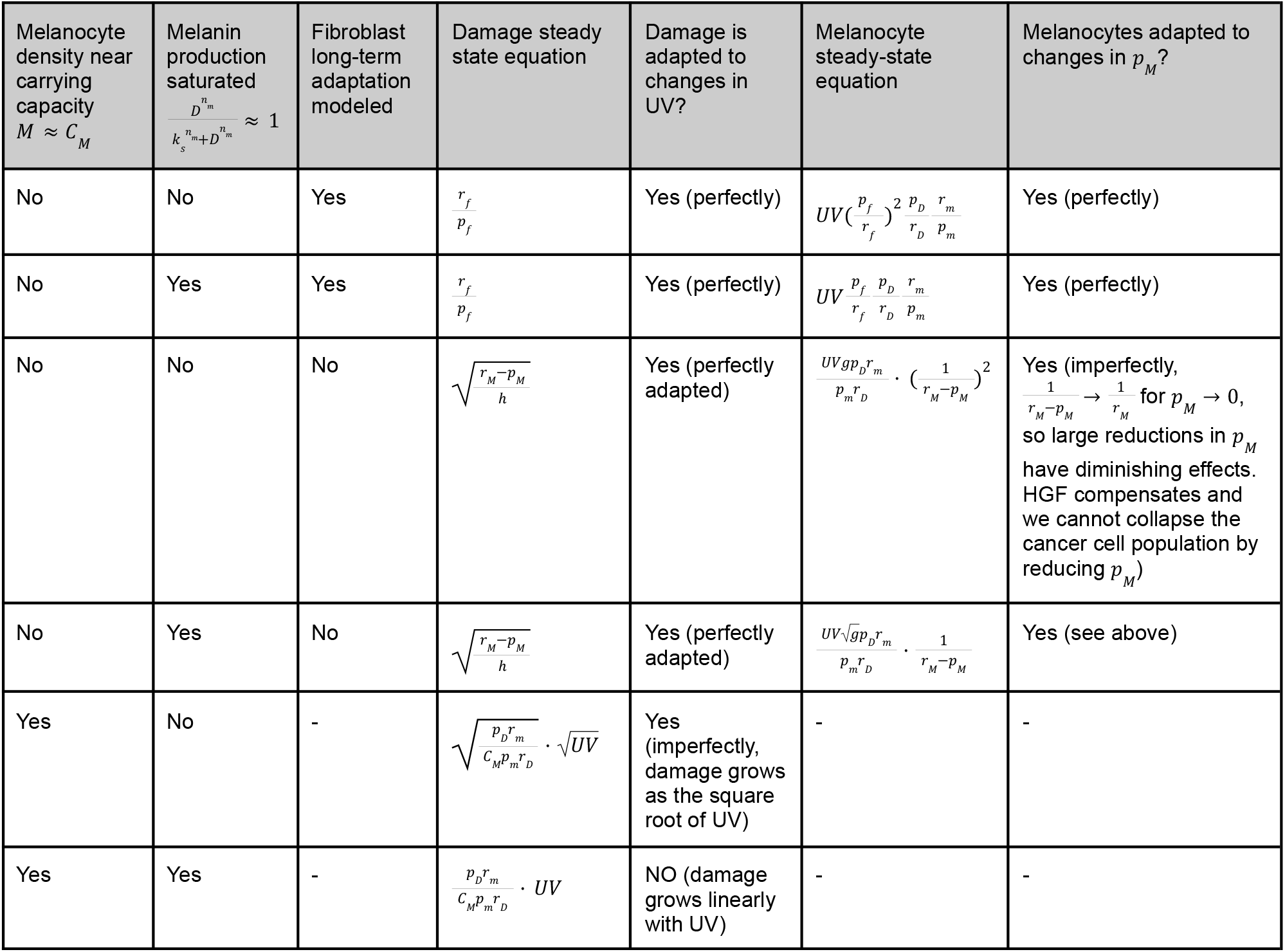
Analytical formulas for UV damage and melanocyte steady states under various model assumptions.

**Figure S4.**
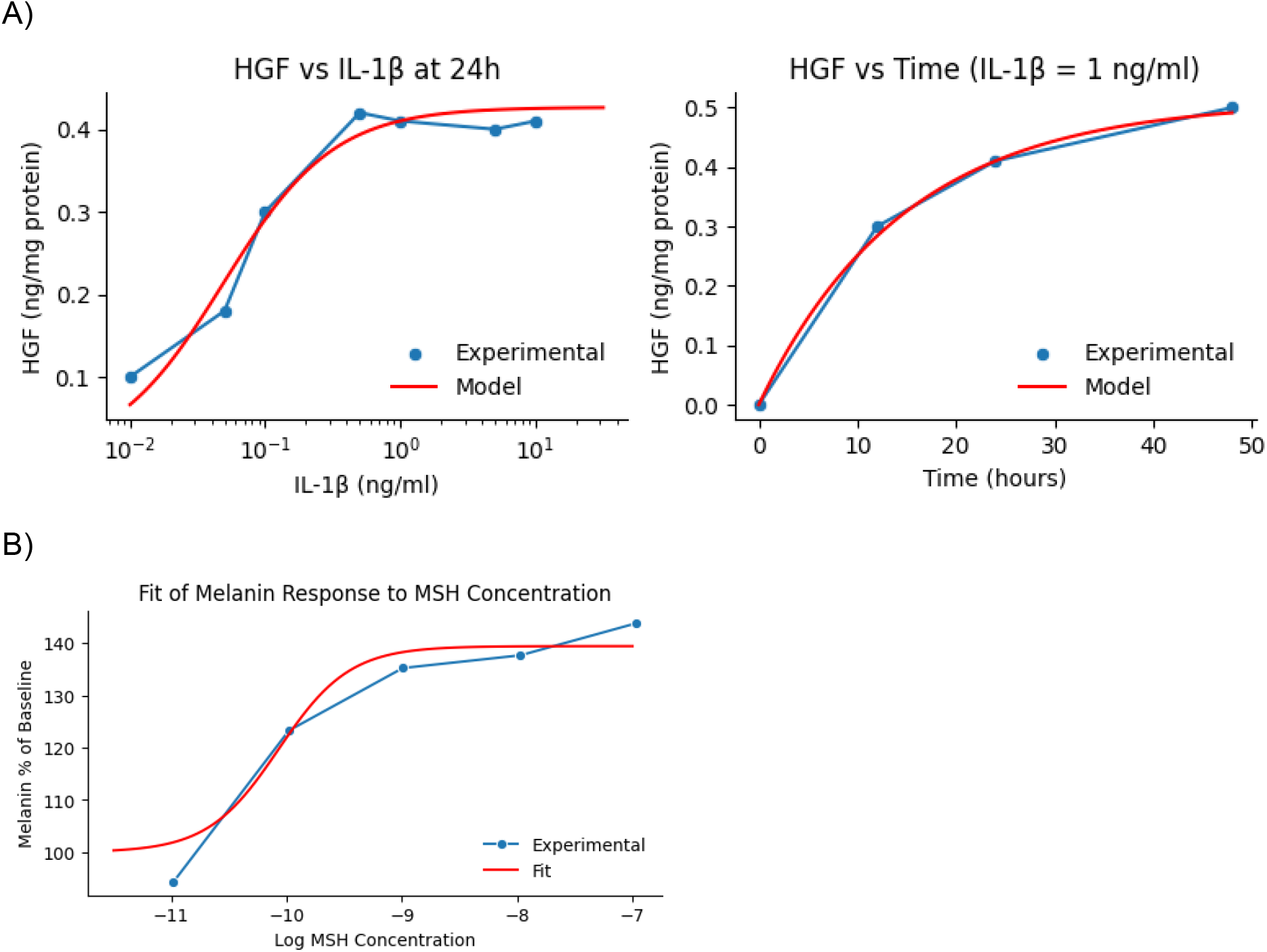
Inferring *n*_*m*_, *n*_*H*_ based on experimental data. A) Matsumoto et. al36 exposed fibroblasts to IL-1 and measured the response to different concentrations (left) and the response over time (right). We fit the parameters of the model 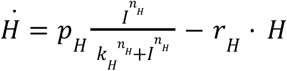. A hill function power of *n*_*H*_ = 1. 06 best fits the data (minimal R^2^). B) Hunt et. al45 quantified melanocyte production of melanin by culturing melanocytes over 3-days with increasing concentrations of MSH. We assume a fixed number of melanocytes over the experiment and a hill-function response to MSH. Isolated melanin is very stable so we neglect its degradation. Thus, 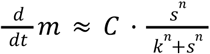, and 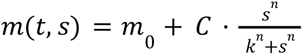. Gillian et. al measured the percentage change in melanin for different concentrations of *s*. We model it as: 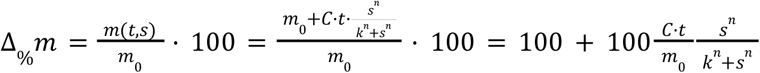. We define *C*′ = 100 · *CT*/*m*_0_, so 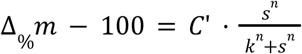. This model has 3 parameters, *C*′, *k, n*. We inferred *n* = 1. 4. In the main section and our analytical derivations of steady states we use *n*_*m*_ = *n*_*H*_ = 1, and in S6,7 we show that results are reproduced for higher values of *n*.

**Table S5.**
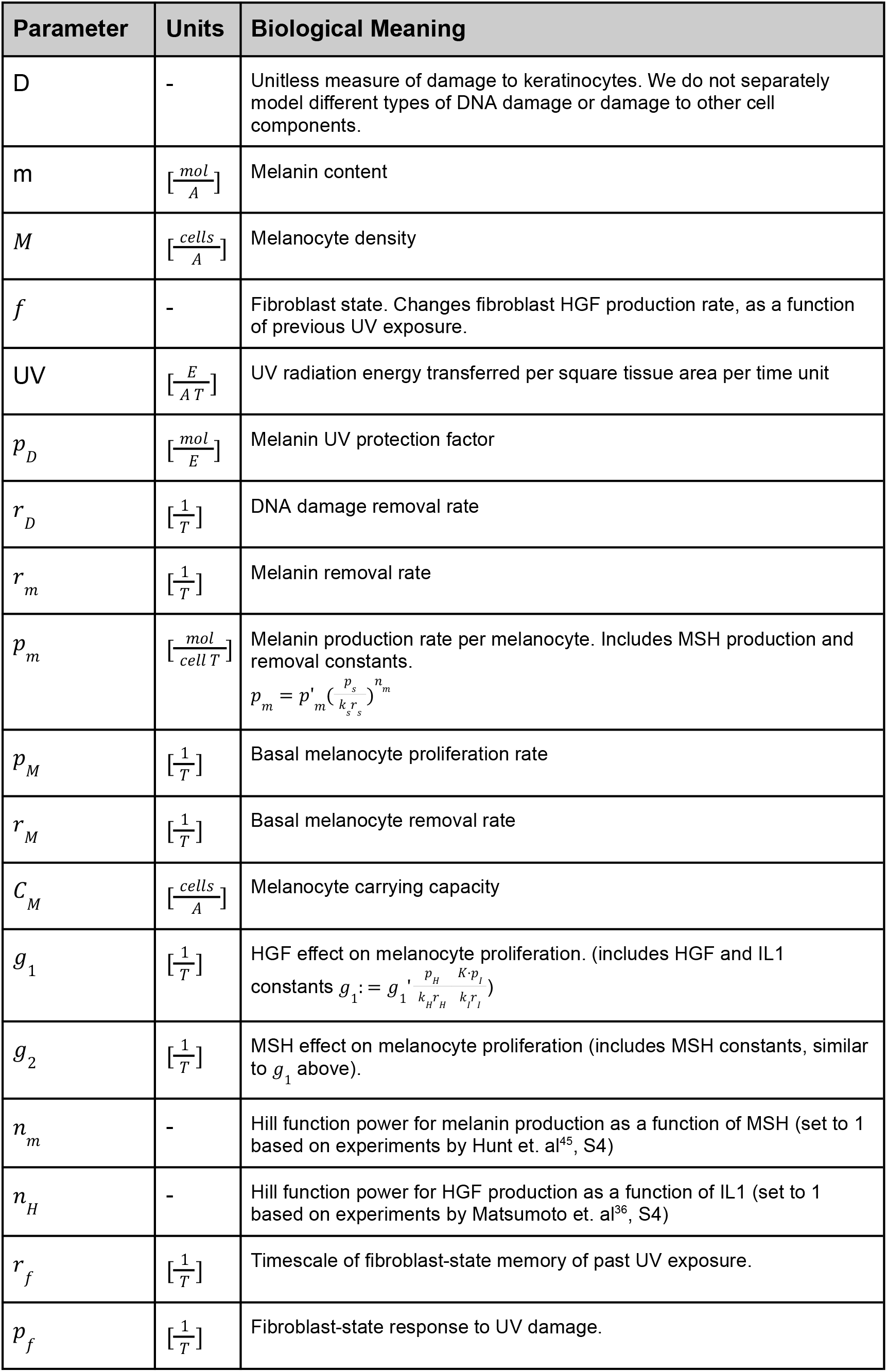
UV homeostasis model parameters and units.

| Parameter | Units | Biological Meaning |
| --- | --- | --- |
| D | - | Unitless measure of damage to keratinocytes. We do not separately model different types of DNA damage or damage to other cell components. |
| m | $[\frac{mol}{A}]$ | Melanin content |
| M | $[\frac{cells}{A}]$ | Melanocyte density |
| f | - | Fibroblast state. Changes fibroblast HGF production rate, as a function of previous UV exposure. |
| UV | $[\frac{E}{AT}]$ | UV radiation energy transferred per square tissue area per time unit |
| $p_D$ | $[\frac{mol}{E}]$ | Melanin UV protection factor |
| $r_D$ | $[\frac{1}{T}]$ | DNA damage removal rate |
| $r_m$ | $[\frac{1}{T}]$ | Melanin removal rate |
| $p_m$ | $[\frac{mol}{cell T}]$ | Melanin production rate per melanocyte. Includes MSH production and removal constants.<br>$p_m = p'_m (\frac{p_s}{k_s r_s})^{n_m}$ |
| $p_M$ | $[\frac{1}{T}]$ | Basal melanocyte proliferation rate |
| $r_M$ | $[\frac{1}{T}]$ | Basal melanocyte removal rate |
| $C_M$ | $[\frac{cells}{A}]$ | Melanocyte carrying capacity |
| $g_1$ | $[\frac{1}{T}]$ | HGF effect on melanocyte proliferation. (includes HGF and IL1 constants $g_1 := g'_1 \frac{p_H}{k_H r_H} \frac{K p_I}{k_I r_I}$) |
| $g_2$ | $[\frac{1}{T}]$ | MSH effect on melanocyte proliferation (includes MSH constants, similar to $g_1$ above). |
| $n_m$ | - | Hill function power for melanin production as a function of MSH (set to 1 based on experiments by Hunt et. al<sup>45</sup>, S4) |
| $n_H$ | - | Hill function power for HGF production as a function of IL1 (set to 1 based on experiments by Matsumoto et. al<sup>36</sup>, S4) |
| $r_f$ | $[\frac{1}{T}]$ | Timescale of fibroblast-state memory of past UV exposure. |
| $p_f$ | $[\frac{1}{T}]$ | Fibroblast-state response to UV damage. |

**Table S6.** Accuracy of tissue classification based on cancer expression of ligands or receptors. We fit logistic regression models to ligand or receptor gene expression data from normal samples, with the aim of detecting the organ an epithelial cell is from (Methods). The model was fit only to data from healthy tissue samples. We apply the model to 350 tumor samples from all organ types. Table S6 shows the area under the receiver operating characteristic curves for models fit separately to ligand or receptor data. In each setting, a total of 350 tumor samples were classified (including all organs). The rightmost column shows the number of samples from the corresponding organ (i.e the positive samples).

| organ | Receptor or ligand gene expression | AUC | Number of positive samples (total n=350) |
| --- | --- | --- | --- |
| breast | ligand | 0.913 | 96 |
| breast | receptor | 0.91 | 96 |
| colorectal | ligand | 0.999 | 25 |
| colorectal | receptor | 0.999 | 25 |
| kidney | ligand | 0.988 | 29 |
| kidney | receptor | 0.996 | 29 |
| liver | ligand | 0.999 | 21 |
| liver | receptor | 1.0 | 21 |
| lung | ligand | 0.978 | 65 |
| lung | receptor | 0.959 | 65 |
| ovarian | ligand | 0.902 | 56 |
| ovarian | receptor | 0.843 | 56 |
| prostate | ligand | 0.956 | 28 |
| prostate | receptor | 0.93 | 28 |
| skin | ligand | 1.0 | 30 |
| skin | receptor | 0.974 | 30 |

**Table S7.** Cancer types and their corresponding cell types of origin.

| Organ | Malignancy | Cell type of origin |
| --- | --- | --- |
| breast | Breast Cancer | luminal epithelial cell of mammary gland |
| colorectal | Colorectal Cancer | intestinal crypt stem cell of colon |
| kidney | Clear Cell Renal Cell Carcinoma | kidney epithelial cell |
| liver | Hepatocellular Carcinoma | hepatocyte |
| lung | Lung Adenocarcinoma | pulmonary alveolar type 2 cell |
| ovarian | Ovarian Cancer | ovarian surface epithelial cell |
| prostate | Prostate Cancer | luminal cell of prostate epithelium |
| skin | Melanoma | melanocyte |

## Additional material

Table S8 - Cell type definitions (Attached)

Table S9 - Top homeostatic receptors per tissue

Table S10 - Top homeostatic ligands per tissue

